# Phylogenetic reconstruction and divergence time estimation of Blumea DC. (Asteraceae: Inuleae) in China based on nrDNA ITS and cpDNA trnL-F sequences

**DOI:** 10.1101/497073

**Authors:** Ying-bo Zhang, Yuan Yuan, Yu-xin Pang, Fu-lai Yu, Dan Wang, Xuan Hu, Xiao-lu Chen, Ling-liang Guan, Chao Yuan

## Abstract

The genus Blumea DC. is one of the economically most important genera of Inuleae (Asteraceae) in China. It is particularly diverse in south China, where 30 species occur, more than half of which are used as herbal medicine or in the chemical industry. However, little is known about the phylogenetic relationships and molecular evolution of this genus in China. We used nuclear ribosomal DNA (nrDNA) internal transcribed spacer (ITS) and chloroplast DNA (cpDNA) trnL-F sequences to reconstruct the phylogenetic relationship and estimate the divergence time of Blumea DC. in China. Results indicated the genus of Blumea DC. was monophyletic and could be divided into two clades, the two clades were resolved that differ with respect to habitat, morphology, chromosome type and chemical composition, of their members. Furthermore, based on the two root time of Asteraceae, the divergence time of Blumea DC. were compare estimated. Results indicated the most recent common ancestor of Asteraceae maybe originated from 76–66 Ma, and Blumea DC. maybe originated from 72.71-52.42 Ma, and had an explosive expansion during Oligocene and Miocene (30.09-11.44 Ma), two major clades were differentiated during the Palaeocene and Oligocene (66.29-46.72 Ma), also the evidence from paleogeography and paleoclimate also supported Blumea DC. had an differentiation and explosive expansion during Oligocene and Miocene.

## INTRODUCTION

The genus Blumea DC. is one of the largest genera of Inuleae (Asteraceae), and contains approximately 100 specie(Anderberg & Eldenäs, 2007; Bremer & Anderberg, 1994; Randeria, 1960). It is most diverse in Asia, Africa, and Australia, and has more than one main center of diversity in Africa and South Asia (Randeria, 1960). China is one center of diversity of this genus, with 30 species distributed throughout South China, of which there are 5 species are endemic (Shi, Chen, Chen, Lin, Liu, Ge, Gao, Zhu, Liu, Yang, Humphries, Raab-Straube, Gilbert, Nordenstam, Kilian, Brouillet, Illarionova, Hind, Jeffrey, Bayer, Kirschner, Greuter, Anderberg, Semple, Štěpánek, Freire, Martins, Koyama, Kawahara, Vincent, Sukhorukov, Mavrodiev & Gottschlich, 2011; Zhang & Cheng, 1984). Blumea DC. has economic and ecological value in China. More than half of the species belonging to this genus have medical or ethnobotanical value (Jia & Li, 2005); for example, *B. balsamifera* DC. is an economic source for L-borneol or camphor extraction, and is widely cultivated in the Philippines and China (China, 2000; Huang, Di & Zhao, 2001; Perry & Metzger, 1980). *B. megacephala* and *B. riparia* are important sources of the medical materials for “fuxuekang”(China, 2000). Other species such as *B. aromatica, B. formosana*, and *B. densiflora* are used to treat rheumatism, esophagitis, headache, hypertension, etc(Jia & Li, 2005). The taxonomy, plant morphology, and anatomy of the genus Blumea DC. were first described in China by Zhang and Cheng(Zhang & Cheng, 1984), the authors treated six subsections and 30 species. Since the first description of this genus, its phylogenetic analysis and molecular evolution have been vigorously disputed. Over the years, phylogenetic analysis using molecular markers has been widely used to attempt to resolve this dispute. Molecular phylogenetics based on samples from Thailand, Burma, and Vietnam among other countries was used to assess the generic interrelationships and molecular evolution of Blumea DC. The results showed that Blumea is monophyletic, if the genera Blumeopsis and Merritia are included, and suggested this genus could be divided into two main clades, which differ in respect to habitat, ecology, and distribution (Pornpongrungrueng, Borchsenius, Englund, Anderberg & Gustafsson, 2007). In the present study, we have extended the molecular phylogenetic analysis of Blumea DC. by (a) adding several samples from China to provide a more thorough coverage of the overall distribution and diversity of Blumea DC., and (b) performing divergence time estimation. We aimed to (a) estimate the molecular evolution and phylogenetic relationships of Blumea DC. with a focus on the genus in China, and (b) offer the best hypothesis for the divergence time and evolutionary events of this genus in China.

## MATERIALS AND METHODS

### Plant materials

In this study,16 Chinese samples, including 12 species belong to 3 sections of Blumea DC., were new collected and sequenced, and the specimens of those samples were deposited in the herbarium of the Chinese Academy of Tropical Agricultural Sciences (herbarium code: CATCH).. Moreover, 9 reference sample of Blumea DC. and 7 sample of outgroup (7 species, 6 genera) were download from Genbank. The sequence information, locality, and other details of the accessions are given in Table 1.

**Table 1.** Localities and samples used in the present study

| Section | Code | Latin name | Locality | GenBank accession number |  | Reference |
| --- | --- | --- | --- | --- | --- | --- |
|  |  |  |  | ITS | trnL-F |  |
| **Semivestitae**<br>(2 spp.) | *B. megacephala* | *Blumea megacephala* (Randeria) Chang & Tseng | Guangxi, China | KP052666 | KP052682 | This research |
|  | *B. riparia* | *B. riparia* (Bl.) DC. | Yunnan, China | KP052668 | KP052685 | This research |
| **Macrophyllae**<br>(6 spp.) | *B. balsamifera* sp1 | *B. balsamifera* (L.) DC. | Guizhou, China | KP052658 | KP052674 | This research |
|  | *B. balsamifera* sp2 | *B. balsamifera* (L.) DC. | Yunnan, China | KP052659 | KP052675 | This research |
|  | *B. balsamifera* sp3 | *B. balsamifera* (L.) DC. | Hainan, China | KP052660 | KP052676 | This research |
|  | *B. aromatic* sp1 | *B. aromatica* DC. | Guizhou, China | KP052656 | KP052672 | This research |
|  | *B. aromatic* sp2 | *B. aromatica* DC. | Hainan, China | KP052657 | KP052673 | This research |
|  | *B. oxydonta* | *B. oxydonta* DC. | Thailand | EU195665 | EU195630 | Pornpongrungrueng et al. 2009 |
|  | *B. densiflora* | *B. densiflora* DC. | Thailand | EF210934 | EF211029 | Pornpongrungrueng et al. 2007 |
|  | *B. saxatilis* | *B. saxatilis* Zoll. & Mor. | Australia | EF210945 | EF211040 | Kanis 1993 |
|  | *B. formosana* | *B. formosana* Kitam | Zhejiang, China | KP052665 | KP052678 | This research |
| **Paniculatae**<br>( 9 spp.and 2 varieties) | *B. clarkei* | *B. clarkei* Hook. f. | Thailand | EF210974 | EF211069 | Pornpongrungrueng et al. 2007 |
|  | *B. fistulosa* | *B. fistulosa* (Roxb.) Kurz. | Yunnan, China | KP052661 | KP052677 | This research |
|  | *B. hieraciifolia* | *B. hieraciifolia* (D. Don) DC. | Yunnan, China | KP052662 | KP052679 | This research |
|  | *B. hieraciifolia* var. hamiltonii | *B. hieraciifolia* var. hamiltonii | Burma | EF210972 | EF211067 | Pornpongrungrueng et al. 2007 |
|  | *B. hieraciifolia* var. *macrostachya* | *B. hieraciifolia* var. *macrostachya* | Thailand | EF210937 | EF211032 | Pornpongrungrueng et al. 2007 |
|  | *B. lanceolaria* | *B. lanceolaria* (Roxb.) Druce. | Guangxi, China | KP052664 | KP052681 | This research |
|  | *B. lacera* | *B. lacera* (Burm. f.) DC. | Zhejiang, China | KP052663 | KP052680 | This research |
|  | *B. mollis* | *B. mollis* (D. Don) Merr. | Hainan, China | KP052670 | KP052683 | This research |
|  | *B. napifolia* | *B. napifolia* DC. | Thailand | EF210959 | EF211054 | Pornpongrungrueng et al. 2007 |
|  | *B. oblongifolia* | *B. oblongifolia* Kitam | Hainan, China | KP052667 | KP052684 | This research |
|  | *B. saussureoides* | *B. saussureoides* Chang & Tseng | Yunan, China | KP052669 | KP052686 | This research |
|  | *B. sinuata* | *B. sinuata* (Lour.) Merr. | Thailand | EF210948 | EF211043 | Pornpongrungrueng et al. 2007 |
|  | *B. virens* | *B. virens* DC. | Thailand | EF210957 | EF211052 | Pornpongrungrueng et al. 2007 |
| **Uncertainty** | *Blumeposis. flava* | *Blumeposis. flava* Gagnep. | Thailand | EF210960 | EF211055 | Pornpongrungrueng et al. 2007 |
| **Outgroup** | *Caesulia axillaris* | *Caesulia axillaris* Roxb. | India | EF210949 | EF211044 | Pornpongrungrueng et al. 2007 |
|  | *Pluchea carolinensis* | *Pluchea carolinensis* (Jacq.) G. Don. | Taiwan, China | AF437850 | EU385104 | Panero& Funk. 2008 |
|  | *Laggera alata* | *Laggera alata* (D. Don) Sch.-Bip. ex Oliv. | Thailand | EF210930 | EF211025 | Pornpongrungrueng et al. 2007 |
|  | *L. pterodonta* | *L. pterodonta* (DC.) Sch. Bip. ex Oliv. | Thailand | EF210929 | EF211024 | Pornpongrungrueng et al. 2007 |
|  | *Schlechtendalia luzulifolia* | *Schlechtendalia luzulifolia* | Australia | KF989506 | KF989612 | Funk et al. 2014 |
|  | *Barnadesia caryophylla* | *Barnadesia caryophylla* | Australia | AY504686 | AY504768 | Funk et al. 2014 |
|  | *Elephantopus scaber* | *Elephantopus scaber* L. | Hainan, China | KP052671 | KP052687 | This research |

### DNA isolation

Genomic DNA was isolated from the above accessions by using the QIAGEN DNeasy Plant Minikit (QIAGEN, Düsseldorf, Germany), according to the manufacturer’s instructions. The DNA was diluted to 30 ng/μL, and the ultraviolet absorption values at A260 were used.

### DNA amplification and sequencing

ITS and trnL-F sequence amplification and sequencing: Internal transcribed spacer (ITS) and trnL-F sequence amplification and analysis were conducted according to previously established protocols (Gustafsson, Pepper, Albert & Källersjö, 2010; Taberlet, GiellyGuy & Bouvet, 1991). The PCR mixture consisted of 60 ng DNA, 1.0 μM of each primer (Invitrogen Corp., Carlsbad, CA, USA), 25.0 μL of 2× Taq PCR Master Mix (0.1 U/μL Taq polymerase, 500 μM/μL of each dNTP, 20 mM/μL Tris-HCl at pH 8.3, 100 mM/μL KCl, 3 mM/μL MgCl_2_; Tiangen, China) in a 50-μL volume. The PCR cycle protocol was based on previously established methods(Gustafsson et al., 2010; Taberlet et al., 1991). The PCR products obtained were separated on a 1.2% agarose gel, and then, bands of the expected size were excised from the gel and purified using the QIAquick Gel Extraction Kit (QIAGEN Inc., Valencia, CA, USA), according to the manufacturer’s instructions. The purified PCR products were subjected to sequencing (Invitrogen, Shanghai, China).

### Sequence processing, conversion, and analysis

Single sequence: First, the raw sequences of the ITS and trnL-F fragments (Invitrogen, Shanghai, China) were edited to remove the low quality parts. The edited sequences were then submitted to GenBank (http://www.ncbi.nlm.nih.gov). The sequences of both loci were aligned using MEGA 7.0 (https://www.megasoftware.net/) (Kumar, Stecher & Tamura, 2016) with default options, and saved in the *.nexus and *.aln formats. The interleaved *.nexus files were converted to the non-interleaved *.nex format by using PAUP 4.0a164 (https://paup.phylosolutions.com/)(Swofford, 2003). The sequence saturation and best substitution model were analyzed using DAMBE(http://dambe.bio.uottawa.ca/Include/software.aspx) (Xia & Sudhir, 2018) and jmodeltest 2.1.7 (http://evomics.org/learning/phylogenetics/jmodeltest/)(Darriba, Taboada, Doallo & Posada, 2012), Then the poorly aligned positions and divergent regions in each aligned sequence were removed by Gblocks0.91b (http://molevol.cmima.csic.es/castresana/Gblocks/Gblocks_documentation.html) (Castresana, 2000; Talavera & Castresana, 2007), and the filtration parameters used with were described by Guillou (Talavera & Castresana, 2007).

#### Combined sequence

The datasets of ITS and trnL-F sequences were loaded into PAUP 4.0 (https://paup.phylosolutions.com/) (Swofford, 2003) for compatibility testing (ILD test) (Farris, Källersjö, Kluge & Bult, 2010), the compatibility testing (ILD test) result reveals there is no incongruence between ITS and trnL-F sequences (p=0.19). Then, the sequences were merged using SequenceMatrix1.8 (https://www.softpedia.com/get/Science-CAD/Sequence-Matrix.shtml) (Vaidya, Lohman & Meier, 2011), and saved in the *.nex format.

### Phylogenetic tree construction

In order to reconstruct a more accurate phylogeny of Blumea DC., first, the single sequence or combined sequence were respectively used for construct a phylogeny tree with four phylogenetic tree construction methods, include the Neighbor-joining (NJ) method, Maximum Parsimony (MP) method, Maximum likelihood (ML) method and Bayesian inference (BI) method in PAUP 4.0a164 (https://paup.phylosolutions.com/) (Swofford, 2003), IQ-TREE (http://www.iqtree.org/)(Chernomor, Von & Minh, 2016; Lam-Tung, Schmidt, Arndt & Bui Quang, 2015) and MrBayes 3.2.6 (http://nbisweden.github.io/MrBayes/download.html) (Huelsenbeck & Ronquist, 2001). Then the topology of phylogenetic tree with different methods were evaluated by TreePuzzle 5.3.rc16 (http://www.tree-puzzle.de/)(Schmidt, Petzold, Vingron & Haeseler, 2003) and CONSEL (http://stat.sys.i.kyoto-u.ac.jp/prog/consel/) software (Shimodaira & Hasegawa, 2001). The phylogenetic trees with best topology were edited with FigTree 1.4.2 (http://tree.bio.ed.ac.uk/software/figtree/) and Inkscape (https://inkscape.org/).

#### Neighbor-joining (NJ) method

The NJ analysis with the PAUP 4.0a164 version (Swofford, 2003) was performed with heuristic searches under the random option of stepwise addition algorithm with equal weighting of all characters. Bootstrap 50% majority-rule consensus trees were reconstructed by performing the heuristic searches with 1,000 bootstrap replicates.

#### Maximum Parsimony (MP) method

The same with NJ method, an Maximum Parsimony(MP) analysis were also reconstructed with the PAUP 4.0a164 (Swofford, 2003), and performing the heuristic searches with 1,000 bootstrap replicates.

#### Maximum likelihood (ML) method

Before Maximum likelihood (ML) analysis with IQ-TREE(Chernomor et al., 2016; Lam-Tung et al., 2015). The best substitution models of single or combined sequences were computed with jmodeltest2.1.7(Darriba et al., 2012). Then the Phylogenetic tree with maximum likelihood (ML) method were analyzed with IQ-TREE (Chernomor et al., 2016; Lam-Tung et al., 2015)with default settings, and 1000 bootstrap (BP) values for tree evaluation.

#### Bayesian inference (BI) method

With the best substitution models parameters computed by jmodeltest 2.1.7 (Darriba et al., 2012), Bayesian inference analysis (BI) analysis were performed with MrBayes 3.2.6 (Huelsenbeck & Ronquist, 2001). The analysis parameters were set as four chains ran simultaneously for 10,000,000 generations or until the average standard deviation of split frequencies fell below 0.01. And the trees were being sampled with every 100 generations, a total of 20,000 trees were generated with the initial sample. Subsequently, the consensus tree were summarized with LogCombiner 1.10.4 (http://beast.community/logcombiner) and TreeAnnotator 1.10.4 (http://beast.community/treeannotator), and the parameters were set as first 10% were discarded as burn-in and 50% majority-rule consensus tree with the posterior probability (PP) values.

### Divergence time estimation

#### Time node selection and correction

Because there are no unequivocal fossils for *Blumea* DC. and its relatives in Inuleae, the divergence and diversification times were difficult to estimate. In order to have an accurate divergence time estimation with Blumea DC., in this study, two root ages of Asteraceae were compared, First, reference the recently results by Barreda et al (Barreda, Luis, Tellería, Olivero, J Ian & Félix, 2015), the root age of Asteraceae as 76–66 Ma; Second, reference the most received time by other researchers, the root age of Asteraceae were set as 49–42 Ma (Kim, Choi & Jansen, 2005; TORICES, 2010; Wagstaff, Breitwieser & Ito, 2011). Subsequently, the geological event time of Qiongzhou strait and Hainan island 5.8–3.7 Ma, were selected as the latest differentiation time of *B. aromatica* DC. samples from Guizhou and Hainan(Huang, Guo, Ho, Shi, Li, Li, Cai & Wang, 2013; Zhao, Wang & Yuan, 2007; Zhu, 2016). Subsequently, in order to investigate the speciation rates of Blumea DC. along with time, BAMM (http://bamm-project.org/) (Rabosky, 2014) and BAMMtools (https://cran.rstudio.com/web/packages/BAMMtools/index.html) (Rabosky, Grundler, Anderson, Title, Shi, Brown, Huang & Larson, 2014) were used for compare the speciation rates with the two hypothesis between two root age of Asteraceae.

#### Divergence time estimation and evolutionary event hypothesis

All divergence times and evolutionary events were hypothesized using BEAST 1.10.4 (http://beast.community/) (Suchard, Lemey, Baele, Ayres, Drummond & Rambaut, 2018). First, the Likelihood ratio test (LRT) (Felsenstein, 1988) were used to determine the ITS or trnL-F data were conformed to molecular clock hypothesis. Results indicates the Blumea data weren’t suitable the molecular clock hypothesis (P = 0.03). Then the divergence times were analyzed with BEAST 1.10.4 (Suchard et al., 2018) incorporating with an uncorrelated lognormal clock model and a birth–death speciation process, and the nucleotide substitution model settings were with jmodeltest result. During the analysis process, the parameters were set as the similar parameters with MrBayes 3.2.6 (Huelsenbeck & Ronquist, 2001), 10,000,000 generations for four independent Markov chain Monte Carlo (MCMC) runs, sampling with every 10000 generations, with substitution and clock models unlinked between partitions. Convergence and effective sample size (> 200) of parameters were analyzed with Tracer 1.7 (http://tree.bio.ed.ac.uk/software/tracer/). LogCombiner 1.10.4 (http://beast.community/logcombiner) and TreeAnnotator 1.10.4 (http://beast.community/treeannotator) were used for summarized the trees, and parameters were set as first 10% were discarded as burn-in and 50% majority-rule consensus tree with the posterior probability (PP) values. Last, in order to reflect the divergence time of Blumea DC., especially the evolutionary time with geological time scale, a strap package(Bell & Lloyd, 2015) in R was used for visualization of the results of BEAST.

## RESULTS

### Features of nrDNA ITS and cpDNA trnL-F sequences in *Blumea* DC

Based on the results of sequence features and substitution models of single sequence or combined sequence from MEGA 7.0 (Kumar et al., 2016), DAMBE(Xia & Sudhir, 2018), and jmodeltest 2.1.7 (Darriba et al., 2012) (Table 2.), results indicated: (1) the nrDNA ITS sequence ranged from 694 ~ 738 bp, after aligned the sequence contained 657 characters, including 230 constant characters (35.01%), 314 informative characters (47.79%), 114 uninformative characters (17.20%). (2) The cpDNA trnL-F sequence ranged from 766~850 bp, after aligned the sequence contained 858 characters, including 730 constant characters (85.08%), 58 informative characters (6.76%), 70 uninformative characters (8.16%). (3) And the combined sequence ranged from 1423~1507 bp, including 960 constant characters (63.49%), 370 informative characters (24.47%). And also the results from the compatibility testing (ILD test) with PAUP 4.0a164 (Swofford, 2003) indicated, the nrDNA ITS sequence and cpDNA trnL-F sequence can combined, and with p=0.19.

**Table 2.** Features of nrDNA ITS and cpDNA trnL-F sequences of *Blumea* DC.

| DNA region | Number of characters | Number of constant characters | Number of variable characters | Number of informative characters | Percent of informative Sites (%) | Best substitution model | Model matrix |  |  |  |
| --- | --- | --- | --- | --- | --- | --- | --- | --- | --- | --- |
|  |  |  |  |  |  |  | Rates | Ncat | P-invar | Gamma shape |
| nrDNA ITS | 657 | 230 | 427 | 314 | 47.79 | SYM+I+G | Gamma | 4 | 0.2390 | 2.1770 |
| cpDNA trnL-F | 858 | 700 | 158 | 70 | 8.16 | TVM+G | Gamma | 4 | 0.0 | 0.9240 |
| Combined sequence | 1545 | 960 | 551 | 375 | 24.27 | - | - | - | - | - |

Also the results of best substitution models from jmodeltest 2.1.7 with AIC information criterion (Akaike, 1973) and BIC information criterion (Schwarz, 1978) indicated, the best substitution models of ITS, trnL-F or combined were SYM+I+G, TVM+G, and GTR+I+G (Table 2), other parameters could also find from Table 2.

### Phylogenetic relationship of Blumea DC

Based on the sequence of nrDNA ITS, the results of four different methods all indicated the genus of Blumea DC. was monophyletic, but the topology of four methods were somehow difference. For example, the topology of NJ method (Fig. 1A) indicated the genus of Blumea DC. could be divided into two clades, the phylogenetic relationship of *B. aromatica* clade maybe clear, but the phylogenetic relationship of others were disordered, also with very low posterior probabilities. The results from MP method (Fig. 1B) seem to mostly conform to the relationship of biological property in Blumea DC., the genus could be divided into three clades, clade B. balsamifera was the basal clade of this genus, the *B. aromatica* clade maybe the transitional clade of this genus, and others maybe the higher group, also with more extensive environmental adaptability, such as with one year herbal, diverse habitat, and others. The results with BI method (Fig. 1C.) seem to have same results with NJ method, also with high posterior probabilities, but pitched the *Caesulia axillaris* into *B. aromatica* clade. Also the topology with different methods were compared with TreePuzzle 5.3.rc16 (http://www.tree-puzzle.de/) (Schmidt et al., 2003) and CONSEL (http://stat.sys.i.kyoto-u.ac.jp/prog/consel/) (Shimodaira & Hasegawa, 2001), results indicated ML method were the most suitable tree-reconstructed method for ITS sequence, and with first Rank (Table 3). Based on the phylogenetic tree with ITS sequence and MP method (Fig. 1D), the genus of Blumea DC. could be divided into two major clades. Clade I were consisted with three species of Macrophyllae (including *B. aromatica, B. densiflora* and *B. balsamifera*) and one species of section Paniculatae (*B. lanceolaria*), but with very low bootstrap. Clade II, with a very perfect bootstrap, was consisted with three subclades and two single species, *B. virens* and *B. fistulosa*. Subclade I were consisted with three species of section Paniculatae (including *B. saussureoides, B. sinuata, B. oblongifolia*, and *B. lacera*), also very perfect bootstrap value. Subclade II were consisted with three species (*B. oxyodonta, B. mollis, B. hieraciifolia*) and two variants of B. hieraciifolia (*B. hieraciifolia* var. hamiltoni and *B. hieraciifolia* var. macrostachy). Subclade III were consisted with seven species, but with diverse inscape, including with two species of Semivestitae (*B. megacephala* and *B. riparia*), two species of Macrophyllae (*B. formosana* and *B. saxatilis*), two of Semivestitae (*B. napifolia* and *B. clarkei*), and also *Blumeopsis flava* also belong to this subclade.

**Fig. 1.**
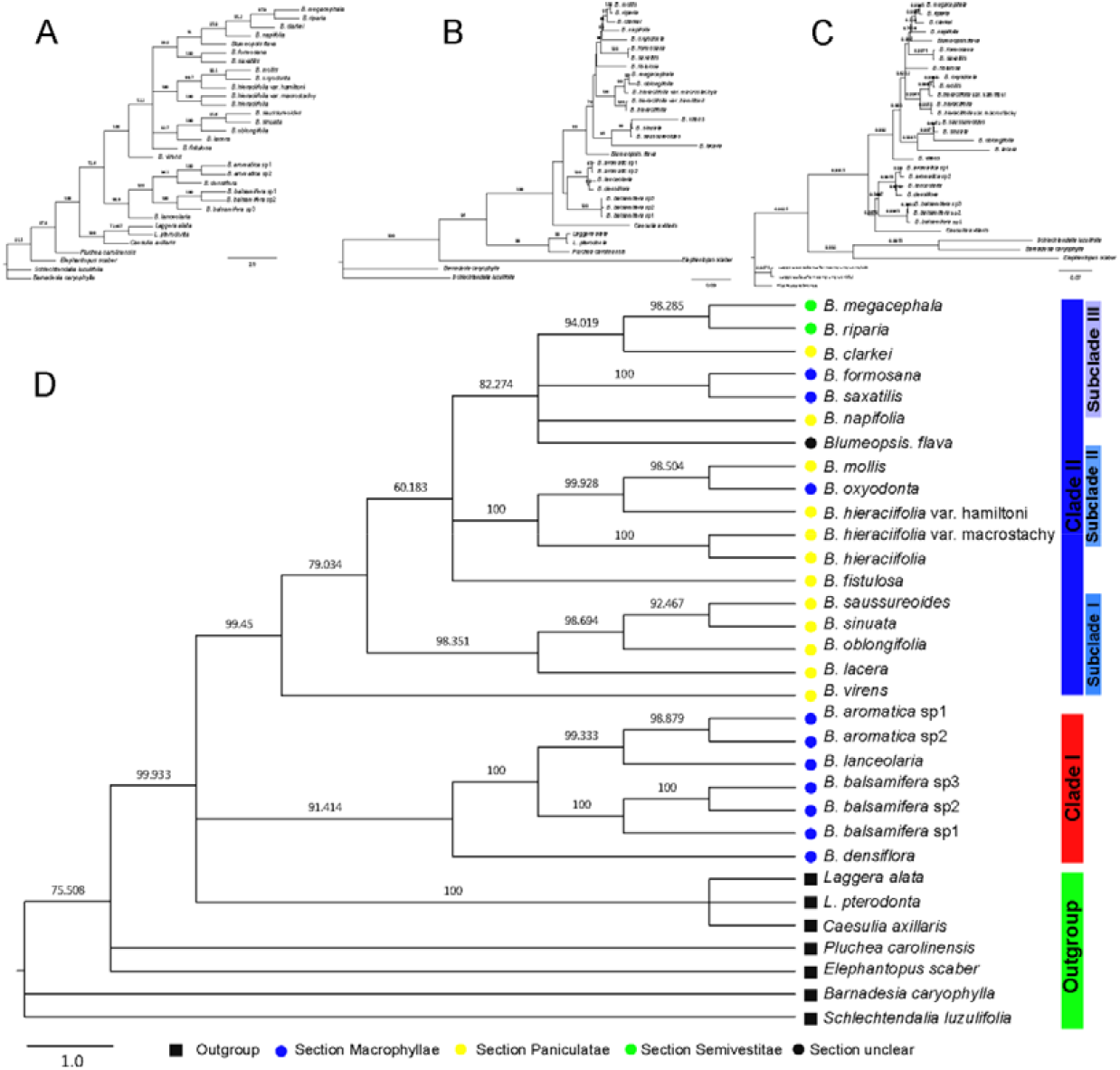
Phylogenetic tree based on the nrDNA ITS sequence, and numbers above branches indicate bootstrap value or posterior probabilities. (A) Phylogenetic tree determined by NJ method with using PAUP program, and numbers above branches indicate bootstrap value. (B) The phylogenetic tree with best topology, determined by MP method with using PAUP program, and numbers above branches indicate bootstrap value. (C) Phylogenetic tree determined by BI method with using MrBayes program, and numbers above branches indicate posterior probabilities. (D) Phylogenetic tree determined by ML method with using IQtree program, and numbers above branches indicate bootstrap value.

**Table 3.** The topology compare with four phylogenetic tree construction method

| DNA region | Tree method | Rank | obs<sup>a</sup> | au<sup>b</sup> | np<sup>c</sup> | bp<sup>d</sup> | pp<sup>e</sup> | kh<sup>f</sup> | sh<sup>g</sup> | wkh<sup>h</sup> | wsh<sup>i</sup> |
| --- | --- | --- | --- | --- | --- | --- | --- | --- | --- | --- | --- |
| nrDNA ITS | NJ | 3 | 2.6 | 0.358 | 0.331 | 0.095 | 0.067 | 0.337 | 0.557 | 0.285 | 0.635 |
|  | MP | 4 | 26.5 | 0.002 | 0.002 | 0.002 | 3E-012 | 0.013 | 0.025 | 0.013 | 0.019 |
|  | ML | 1 | -2.6 | 0.681 | 0.669 | 0.674 | 0.866 | 0.662 | 0.859 | 0.662 | 0.885 |
|  | BI | 2 | 2.6 | 0.357 | 0.331 | 0.229 | 0.067 | 0.338 | 0.558 | 0.338 | 0.648 |
| cpDNA trnL-F | NJ | 4 | 0.4 | 0.337 | 0.274 | 0.275 | 0.176 | 0.300 | 0.309 | 0.300 | 0.309 |
|  | MP | 3 | 0.0 | 0.410 | 0.213 | 0.210 | 0.274 | 0.326 | 0.743 | 0.326 | 0.620 |
|  | ML | 2 | 0.0 | 0.503 | 0.357 | 0.351 | 0.274 | 0.453 | 0.682 | 0.453 | 0.655 |
|  | BI | 1 | -0.0 | 0.731 | 0.163 | 0.164 | 0.275 | 0.547 | 0.997 | 0.547 | 0.992 |
| Combined sequence | NJ | 4 | 8.5 | 0.097 | 0.077 | 0.077 | 1E-004 | 0.125 | 0.229 | 0.125 | 0.187 |
|  | MP | 3 | 0.0 | 0.328 | 0.293 | 0.301 | 0.009 | 0.297 | 0.435 | 0.297 | 0.413 |
|  | ML | 2 | 0.0 | 0.165 | 0.055 | 0.056 | 0.495 | 0.085 | 0.753 | 0.085 | 0.691 |
|  | BI | 1 | -0.0 | 0.867 | 0.578 | 0.566 | 0.496 | 0.915 | 0.982 | 0.915 | 0.986 |

Notes: Obs<sup>a</sup> = observed log-likelihood difference to the best topology; au<sup>b</sup> = approximately unbiased; np<sup>c</sup> = bootstrap probability of the topology (i.e., the probability that the given topology has the largest likelihood in 10 scaled sets of 10,000 bootstrap replicates); bp<sup>d</sup> = np with 10 non-scaled sets of 10,000 bootstrap replicates; pp<sup>e</sup> = Bayesian posterior probabilities of the model; kh<sup>f</sup> = Kishino-Hasegawa; sh<sup>g</sup> = Shimodeira-Hasegawa; wkh<sup>h</sup> = weighted KH; wsh<sup>i</sup> = weighted SH.

With cpDNA trnL-F sequence, the results of four different methods were also indicated the genus of Blumea DC. were also divided into two clades, but the relationship of four methods were somehow difference, especially the species in clade II (Fig2.). Combined the results from topology compare indicated Bayesian inference (BI) method maybe was the most suitable tree-reconstructed method (Table 3). With BI method, Blumea DC. also could been divided into two clades, clade I, also with the same members in nrDNA ITS data set, was consisted with three species of Macrophyllae (including *B. aromatica, B. densiflora* and *B. balsamifera*) and one species of section Paniculatae (*B. lanceolaria*), and the other species were combined into Clade II, but with low posterior probabilities (Fig.2D). Also the same results were observed from NJ, MP and ME method (Fig.2A, Fig.2B and Fig.2C), the genus could be divided into two clades, but with low bootstrap value, especially in the phylogenetic tree in NJ method and MP method, and this may be caused by the little informative characters in trnL-F data set.

**Fig. 2.**
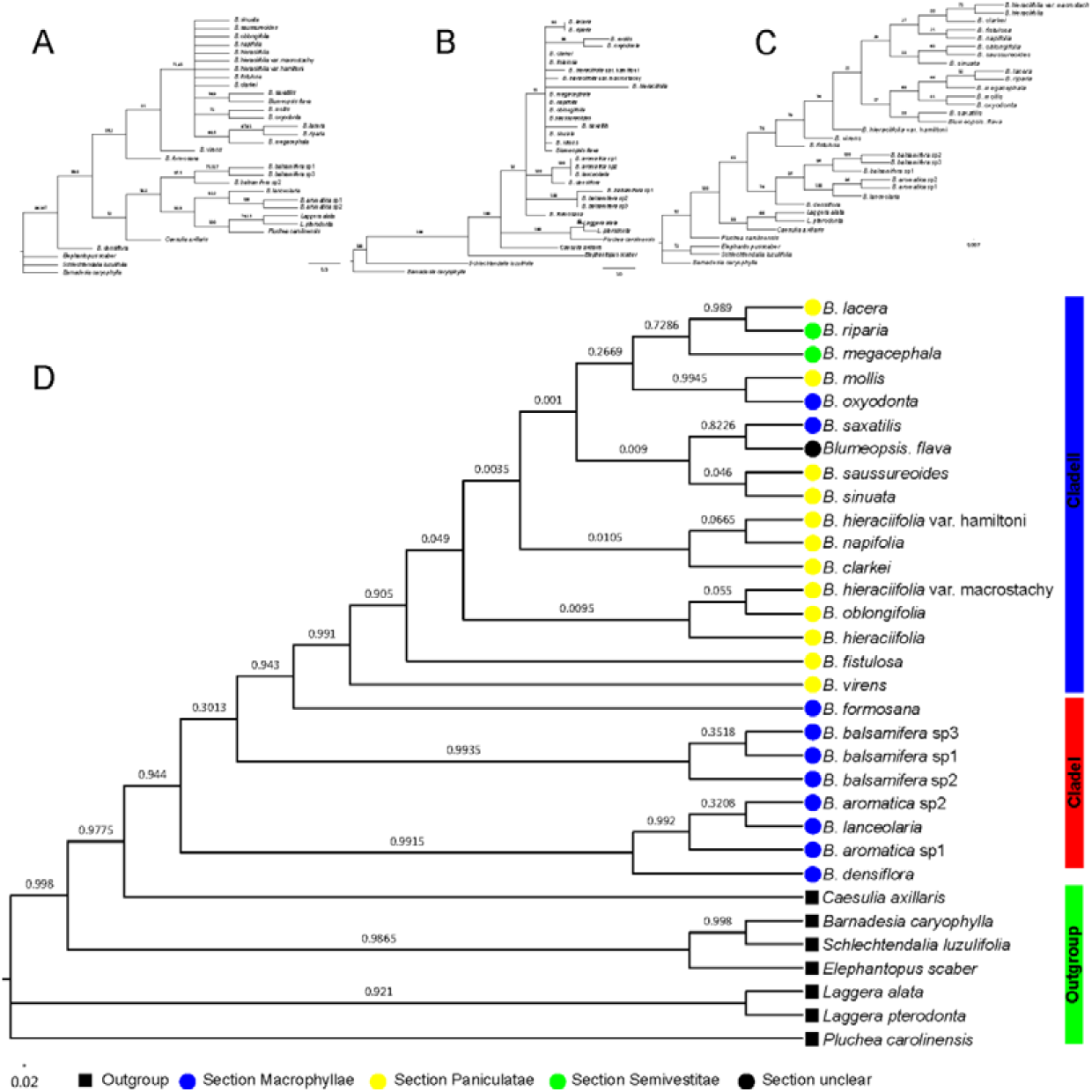
Phylogenetic tree based on the cpDNA trnL-F sequence, and numbers above branches indicate bootstrap value or posterior probabilities. Phylogenetic tree determined by NJ method with using PAUP program, and numbers above branches indicate bootstrap value. (B) The phylogenetic tree with best topology, determined by MP method with using PAUP program, and numbers above branches indicate bootstrap value. (C) Phylogenetic tree determined by ML method with using IQtree program, and numbers above branches indicate bootstrap value. (D) Phylogenetic tree determined by BI method with using MrBayes program, and numbers above branches indicate posterior probabilities.

Furthermore, the combined sequences were also analyzed with both four methods, results indicated Bayesian inference (BI) method was also the most suitable tree-reconstructed method for combined sequences (Table 3), also with the same topology based on the nrDNA ITS sequence (Fig 3.). And the genus of Blumea DC. could be divided into two clades, and clade II was also consisted with three subclades and two single species(Fig 3.D). From the results from NJ, MP and ME method (Fig.3A, Fig.3B and Fig.3C), the same results were observed, and also with better bootstrap posterior supported value. This means the combined sequence can improve the lower posterior probabilities or bootstrap caused by single sequence, also can solve the dispute caused by single sequence.

**Fig. 3.**
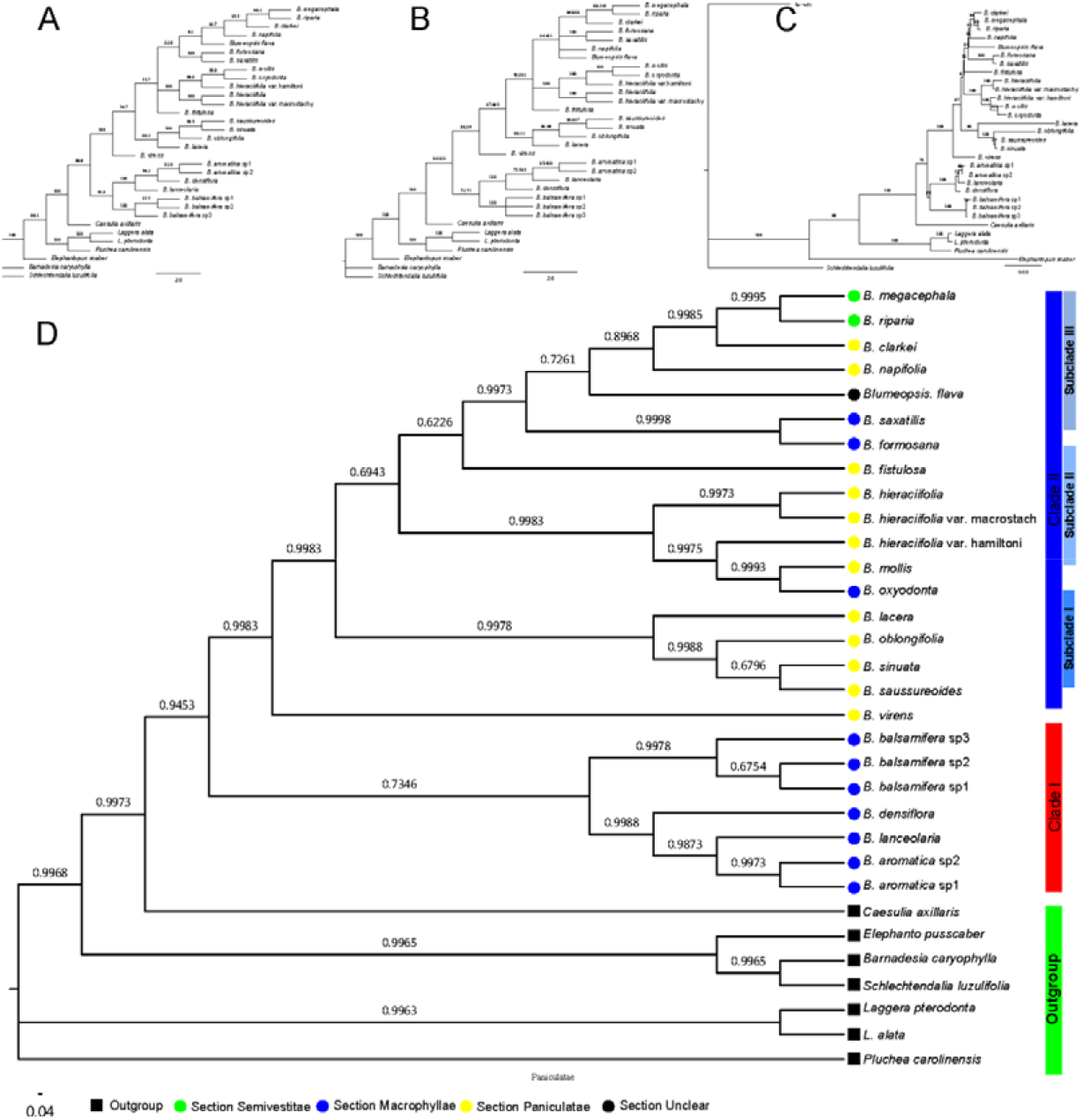
Phylogenetic tree based on the combined sequence, and numbers above branches indicate bootstrap value or posterior probabilities. Phylogenetic tree determined by NJ method with using PAUP program, and numbers above branches indicate bootstrap value. (B) Phylogenetic tree determined by MP method with using PAUP program, and numbers above branches indicate bootstrap value. (C) Phylogenetic tree determined by ML method with using IQtree program, and numbers above branches indicate bootstrap value.(D) Phylogenetic tree determined by BI method with using MrBayes program, and numbers above branches indicate posterior probabilities.

### Divergence time estimation and evolutionary event hypothesis

Based on the compare of two calibration points of root age of Asteraceae and geography-species differentiation time, a series of evolutionary events and divergence times were estimated (Fig. 4). With the root age of Asteraceae as 49–42 Ma (Kim et al., 2005; TORICES, 2010; Wagstaff et al., 2011) (Fig. 4B), the time of genus Blumea DC. differentiated from Inuleae was estimated to be 45.16-33.73 Ma (the time between late of Eocene and early of Oligocene). The divergence time of two major clades in Blumea DC. was estimated to be 42.40-29.28 Ma. During the Oligocene and Miocene, this genus had an explosive expansion, and many species were speciated during the period of 20.05-5.75 Ma. With the root age of Asteraceae as76–66 Ma (Barreda et al., 2015) (Fig. 4D), the time of genus Blumea DC. differentiated from Inuleae was estimated to be 72.71-52.42 Ma (the time between late of Upper and early of Paleocene), the divergence time of two major clades in Blumea DC. was further be estimated to be 66.29-46.72Ma. During Oligocene and Miocene (30.09-11.44 Ma), this genus had an explosive expansion, and many species were speciated during this time. Furthermore, the speciation rate were also estimated with BAMM (Rabosky, 2014) and BAMMtools (Rabosky et al., 2014), and results indicated the speciation rate were higher in the early time(Fig. 4A and Fig. 4D). Combined the speciation rate estimation results and the compare divergence time estimation results with two calibration points of root age of Asteraceae, I believed the with the root age of Asteraceae as76–66 Ma may be had an relative accuracy divergence time estimation for Blumea DC.. First, based on the paleogeography and paleoclimate of Oligocene, the temperature decline in Paleogene Period is interrupted by an Oligocene (~33.5 Ma), and an stepwise climate increase began 32.5 Ma and lasted through to 25.5 Ma(Lisiecki & Raymo, 2005; Miller, Browning, Aubry, Wade, Katz, Kulpecz & Wright, 2008), also the same time is the most important period for the grasslands expanded and forests extent, and the sunflower family had an explosive expansion during this time (Barreda et al., 2015). Second, the early of Miocene (~20 Ma), is the time of formation and differentiation time of Malaysian flora (Smedmark & Anderberg, 2007). In the old Chinese mainland, because of the relative warm climate, was the most vigorous area of the Earth, and numerous mammalian fossils have been found in China (for example in Gansu Province and Qinghai Province).

**Fig. 4.**
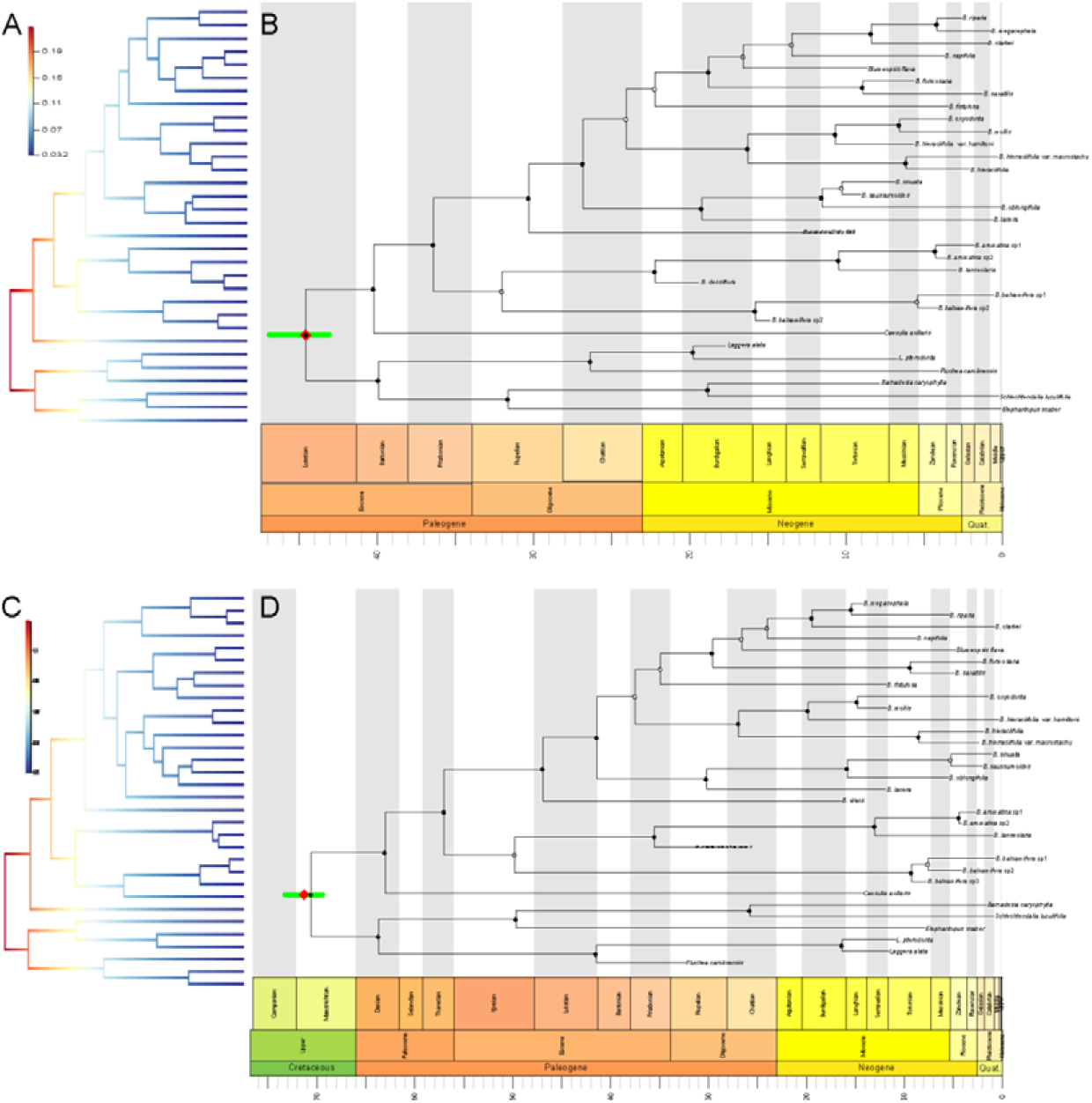
Divergence time estimation and speciation rate analysis based on the combined sequence and unlike model. (A)The speciation rate analysis performed with the root age of Asteraceae as 49–42 Ma. (B) The divergence time estimation phylogenetic tree with geological time scale, and with the root age of Asteraceae as 49–42 Ma, The internal nodes of the tree are indicated with circles, where circles mark nodes with posterior probability:•>0.95, 0.95 >•>0.75, 0.75 >◦. (C)The speciation rate analysis performed with the root age of Asteraceae as 76–66 Ma. (D) The divergence time estimation phylogenetic tree with geological time scale, and with the root age of Asteraceae as 76–66 Ma, The internal nodes of the tree are indicated with circles, where circles mark nodes with posterior probability:•>0.95, 0.95 >•>0.75, 0.75 >◦.

With the evidence from paleogeography and paleoclimate, the time of genus Blumea DC. maybe estimated to be 72.71-52.42 Ma, the divergence time of two major clades was further be estimated to 66.29-46.72Ma, and during Oligocene and Miocene (30.09-11.44 Ma), this genus had an explosive expansion.

## DISCUSSION

The genus Blumea DC. is one of the largest genera of Inuleae (Asteraceae), widely distribute ancient tropical region of around the Pacific, and South China is one center of diversity of this genus, contains 30 species. Because of the variety of its morphology, since the first description of this genus, its phylogenetic analysis and molecular evolution have been vigorously disputed. Over the years, phylogenetic analysis using molecular markers has been widely used to attempt to resolve this dispute (Pornpongrungrueng et al., 2007; Pornpongrungrueng, Borchsenius & Gustafsson, 2009), the results showed that Blumea(included *Blumeopsis flava*) is monophyletic, and suggested this genus could be divided into two main clades and one single species, *B. balsamifera*, which differ in terms of habitat, ecology, and distribution (Pornpongrungrueng et al., 2007). In this study, both nrDNA ITS and cpDNA trnL-F sequences were used to estimate the phylogenetic relationship between the members of this genus. And results indicated this genus could divided into two clades, Clade I included with *B. aromatica*, *B. densiflora*, *B. balsamifera* and *B. lanceolaria*, the others belong to the Clade II. Compare with habit, distribution, chromosome number, chemical composition and etc., we found the two main clades differ in habit, ecology, distribution, chromosome number and chemical composition, the Blumea DC. clade contains mostly perennial shrubs or subshrubs, and mostly distributed from evergreen forests, and also with the same chromosome number (2n = 18)and chromosome structure (6M+8m+4sm) (Lu, 1996), also the chemical composition are similar (Pang, Wang, Fan, Chen, Yu, Hu, Wang & Yuan, 2014). And clade II, which is a widespread paleotropical group that comprises mostly annual, weedy herbs of open forests and fields.

Divergence time estimation and evolutionary event hypothesis is an important content of phylogenetic, but because of the fossil record, it is very difficult to estimate the evolutionary event and divergence time. Choose some geography events as calibration points, could provide more evidence for the process of phylogenetic evolution and divergence event hypothesis (Lanyon & Omland, 1999). But because the fossil record information of Inuleae is very little, most of the families have no fossil record, and several fossils clearly identifiable as members of the Asteraceae and Goodeniaceae from Oligocene and later and seeds of the Menyanthaceae and Campanulaceae from Oligocene and Miocene, respectively (Benton, 1993). There are also several records of Eocene pollen of Asteraceae were found from South America or china (Graham, 1996; Zhu, Wu & Xi, 1985). Because Oligocene pollen of the Asteraceae is of a comparatively specialized type and is found on several continents, it is reasonable to assume that the root time dates back at least to the Oligocene-Eocene boundary 42–49 Ma. Also the same time consulted by Hind, et al. (Hind, Jeffrey & Pope, 1995). But according to a more recent paper by Barreda et al.(Barreda et al., 2015), the most recent common ancestor of Asteraceae maybe originated from 76–66 Ma. In this research, combined the form time of Hainan island and Qiongzhou strait (5.8-3.7 Ma)(Huang et al., 2013; Zhao et al., 2007; Zhu, 2016), the divergence time of Blumea DC. were compare estimated, and results indicated Blumea DC. maybe originated from 72.71-52.42 Ma, and had an explosive expansion during Oligocene and Miocene (30.09-11.44 Ma), two major clades were differentiated during the Paleocene and Oligocene (66.29-46.72 Ma), also the evidence from paleogeography and paleoclimate also supported Blumea DC. had an differentiation and explosive expansion during Oligocene and Miocene (Lisiecki & Raymo, 2005; Miller et al., 2008).

## Acknowledgments

This work was supported by grants from National Natural Science Foundation of China (No. 81374065 and No. 81403035), Modernization of Chinese Traditional Medicines in Hainan province [No. ZY201410], and Basal Research Fund of Central Public-interest Scientific Institution (No. 1630032015020).

